# Intra-individual variability of sleep and nocturnal cardiac autonomic activity in elite female soccer players during an international tournament

**DOI:** 10.1101/664474

**Authors:** Júlio A. Costa, Pedro Figueiredo, Fábio Y. Nakamura, Vincenzo Rago, António Rebelo, João Brito

**Affiliations:** Centre of Research, Education, Innovation and Intervention in Sport, Faculty of Sport, University of Porto, CIFI^2^D, Porto, Portugal; Portugal Football School, Portuguese Football Federation, FPF, Oeiras, Portugal; Research Center in Sports Sciences, Health Sciences and Human Development, CIDESD, University Institute of Maia, ISMAI, Maia, Portugal; Associate Graduate Program in Physical Education UPE/UFPB, João Pessoa, PB, Brazil

**Keywords:** overnight measurements, parasympathetic system, sleep accelerometer, recovery, women football

## Abstract

**Purpose:** This study provides insights into the individual sleep patterns and nocturnal cardiac autonomic activity responses of elite female soccer players during an international tournament.

**Materials and methods:** Twenty elite female soccer players (aged 25.2±3.1 years) wore wrist actigraph units and heart rate (HR) monitors during night-sleep throughout 9 consecutive days (6 day-time training sessions [DT], 2 day-time matches [DM], and 1 evening-time match [EM]) of an international tournament. Training and match loads were monitored using the session-rating of perceived exertion (s-RPE) and wearable 18-Hz GPS (total distance covered [TD], training and match exposure time, and high-speed running [HSR]) to characterize training and match loads.

**Results:** Individually, s-RPE, TD, exposure time, and HSR during training sessions ranged from 20 to 680 arbitrary units (AU), 892 to 5176 m, 20 to 76 min, and 80 to 1140 m, respectively. During matches, s-RPE, TD, exposure time, and HSR ranged from 149 to 876 AU, 2236 to 11210 m, 20 to 98 min, and 629 to 3213 m, respectively. Individually, players slept less than recommended (<7 hours) on several days of the tournament, especially after EM (n=8; TST ranging between 6:00-6:54 h). Total sleep time coefficient of variation (CV) ranged between 3.1 and 18.7%. However, all players presented good sleep quality (i.e., sleep efficiency ≥75%; individual range between: 75-98%) on each day of the tournament. Most of the players presented small fluctuations in nocturnal cardiac autonomic activity (individual nocturnal heart rate variability [HRV] ranged from 3.91-5.37 ms and HRV CV ranged from 2.8-9.0%), while two players presented higher HRV CV (11.5 and 11.7%; respectively).

**Conclusion:** Overall, elite female soccer players seemed to be highly resilient to training and match schedules and loads during a 9 day international tournament.

## Introduction

### Paragraph 1

Elite soccer players are constantly exposed to multiple high physiological demands due to an elevated number of training sessions and matches played in National and international competitions, often with congested match calendars [1]. In some women’s competitive tournaments, only one to two days of recovery are given between matches [2]. In this scenario, optimizing recovery is required to reduce the risk of transitioning into a state of excessive fatigue as well as to reduce the risk of injury [3].

### Paragraph 2

One of the most critical aspects of the recovery *continuum* for elite athletes is obtaining a sufficient quantity and quality of sleep [4]. In fact, athletes and coaches from several sports including soccer have ranked sleep as the most important recovery strategy [5]. A minimum of 7-9 hours of total sleep time (TST) per night is generally recommended to promote optimal health and cognitive function among adults aged 18 to 60 years old [6]. Although there is no general consensus regarding the amount of sleep an elite athlete should obtain to maintain optimal performance [7], athletes who obtain less than 7 hours of sleep per night might have an increased likelihood of injury [4, 8]. In fact, some studies have found sleep durations of less than 7 hours in elite athletes, especially in soccer teams. Sargent et al. [7], for example, found that athletes (from individual and team sports) obtained an average of 6.5 hours sleep per night, ranging from 5 to 8 hours. Lastella et al. [9] confirmed these results, finding that average sleep duration for elite athletes (including elite soccer players) was 6.8 hours, ranging from 5.5 hours to 8 hours. Based on these results, it seems that athletes are probably not getting sufficient sleep. Therefore, and importantly, athletes that might be sleeping for less than 7 hours [10, 11] may require an extension of sleep time. In fact, extended TST (commonly used as sleep quantity index [4]) can lead to better psychomotor accomplishment and technical accuracy [12], with likely positive effects on competitive performance [13]. Besides sleep quantity analyses, sleep efficiency (SE) is recommended for monitoring sleep patterns as a sleep quality variable [4, 14], especially in elite athletes [15]. According to the National Sleep Foundation report [16], SE ≥85% is generally recommended as an appropriate indicator of good sleep quality, whereas a sleep efficiency ≤74% indicates inappropriate sleep quality for young adults/adults. As already mentioned, athletes are often unable to achieve ≥7 hours of TST and ≥85% SE during training and competition [4]. However, these results are especially concerning when interpreted as group mean, suggesting that athletes may achieve these recommendations, where and more likely are included individuals and/or nights that do not achieved [4].

### Paragraph 3

Due to a variety of essential immunological and metabolic processes which occur during sleep, it seems that a conceptual relationship exists between the quantity and quality of sleep and the capacity of athletes to recover and perform [17]. However, the majority of research available examining the sleep of athletes, especially in women, has typically averaged data across several nights, providing a mean estimate of usual sleep [7, 9, 18–20]. While such an approach is useful to provide basic insight into sleep in athletes, it lacks details of how sleep may vary across multiple nights [21]. Moreover, individual variability can reflect differences within individuals over time [22], with high intra-individual sleep variability indicating the need for individualized sleep education strategies and interventions to promote appropriate sleep [21]. Additionally, the coefficient of variation (CV) of sleep parameters (e.g. TST_CV_), classified as a measure of intra-individual sleep variability [23], has been calculated to measure nocturnal sleep variability [24]. In this respect, it may be important to include the presentation of sleep data by encompassing individual responses, in addition to general group means [21]. In addition, special attention should be given to the sleep behavior of elite athletes (e.g., TST_CV_) during international tournaments (a period of highly congested fixtures) since sleep deficits can impair performance [25].

### Paragraph 4

Although sleep is considered a restorative behavior, heart rate HR variability (HRV) has become one of the most practical and popular methods to monitor positive and negative training adaptations in athletes [26]. Recently, there has been growing interest in the use of HRV measurements during sleep to evaluate exercise-induced disturbances in allostatic load (i.e., adaptive processes that maintain homeostasis through the production of mediators such as adrenalin, cortisol, and other chemical messengers) [27], and recovery from daily training and other sources of stress [19, 20, 28]. In fact, it is currently accepted that overnight sleep measurements over consecutive days are appropriate for tracking the recovery of HRV following high-intensity exercise [29]. However, standardized training programs within team sport settings have often produced sparse adaptive results, with high responders and low responders often getting lost in averaged data reports [30]. As a consequence, an increased desire for training individualization in team sport settings has given rise to a variety of athlete-monitoring strategies, enabling coaches to better manage fatigue and manipulate training prescription on an individual basis [31].

### Paragraph 5

Vagal indices of HRV, such as the logarithm of the root mean square of successive R-R interval differences (lnRMSSD), reflecting cardiac parasympathetic modulation, are sensitive to fatigue and have been useful in evaluating individual training adaptation in soccer players [32, 33]. Furthermore, the weekly (4-7 days) CV of lnRMSSD (lnRMSSD_CV_) may provide valuable information concerning training-induced perturbations in homeostasis, i.e., can reflect the day-to-day variations in cardiac parasympathetic activity [32, 34, 35]. In general, athletes with a lower lnRMSSD_CV_ are more aerobically fit and seem to cope better with training and match loads [36–38]. Thus, athletes with high TST and SE and lnRMSSD are expected to experience less perturbation in sleep patterns [23] and cardiac autonomic activity [26].

### Paragraph 6

Although data exist on the role of sleep in recovery and on the impact of various interventions (e.g. competitions, time of day for training, training and match loads) on sleep quality/quantity, there is a lack of individual variability sleep analysis during international tournaments, especially in elite female athletes. These investigations may identify potential factors related to disturbed sleep and nocturnal HRV, which could assist in better defining and recommending appropriate sleep hygiene strategies and more adequately managing fatigue, as well as manipulation of training prescription and post-match recovery on an individual basis. Therefore, the aim of this study was to describe sleeping patterns and nocturnal HRV individual profiles of elite female soccer players from a National team during an international tournament.

## Materials & methods

### Subjects

#### Paragraph 7

Twenty elite outfield female soccer players (age: 25.2±3.1 years; height: 167.2±4.8 cm; body mass: 57.8±3.8 kg; mean ± SD) competing in the Portuguese National team volunteered to participate in the study, during an international tournament (Algarve Cup 2018). The study design was carefully explained to the subjects, and written informed consent was obtained. The study followed the Declaration of Helsinki and was approved by the Ethics Committee of the Faculty of Sports, University of Porto (CEFADE 03.2017).

### Study design

#### Paragraph 8

The study followed a descriptive, observational design, highlighting the individual sleep and overnight HRV responses of elite female soccer players during an international competition. Data collection was performed throughout 9 consecutive days (encompassing 6 day-time training sessions; DT [start ranged between 11:00AM–5:30 PM], 2 day-time matches; DM [both started at 3:00 PM], and 1 evening-time match; EM [started at 7:00 PM) of an international tournament (Table 1). Players’ sleeping patterns and nocturnal cardiac autonomic activity were monitored every night throughout the tournament. Players wore wrist actigraph units and heart rate (HR) monitors during night-sleep. Training and match loads were quantified by session-rating of perceived exertion (s-RPE), total distance covered (TD), training and match exposure time (volume), and high-speed running (HSR) to characterize training practices and competition demands during the observation period.

**Table 1.** Data collection during 9 consecutive days of an international tournament in elite female soccer players. Training load (TL) was assessed each day-training (DT), day-match (DM), and evening-match (EM). Cardiac autonomic activity and actigraphy were assessed during night-sleep, and after each training session and match (represented as grey area). Scheduled time and duration of training and matches are also illustrated.

| 174 | Wednesday | Thursday | Friday | Saturday | Sunday | Monday | Tuesday | Wednesday | Thursday |
| --- | --- | --- | --- | --- | --- | --- | --- | --- | --- |
| 175 | TL (RPE, GPS)<br>Cardiac autonomic activity<br>Actigraphy | TL (RPE, GPS)<br>Cardiac autonomic activity<br>Actigraphy | TL (RPE, GPS)<br>Cardiac autonomic activity<br>Actigraphy | TL (RPE, GPS)<br>Cardiac autonomic activity<br>Actigraphy | TL (RPE, GPS)<br>Cardiac autonomic activity<br>Actigraphy | TL (RPE, GPS)<br>Cardiac autonomic activity<br>Actigraphy | TL (RPE, GPS)<br>Cardiac autonomic activity<br>Actigraphy | TL (RPE, GPS)<br>Cardiac autonomic activity<br>Actigraphy | TL (RPE, GPS)<br>Cardiac autonomic activity<br>Actigraphy |
| 176 | **DT1** | **DT2** | **DM1** | **DT3** | **DM2** | **DT4** | **DT5** | **EM3** | **DT6** |
| 177 | 5:30 PM<br>(65 ± 2 min) | 11:30 AM<br>(70 ± 17 min) | 3:00 PM<br>(67 ± 26 min) | 4:30 PM<br>(34 ± 11 min) | 3:00 PM<br>(79 ± 23 min) | 4:00 PM<br>(71 ± 20 min) | 4:00 PM<br>(57 ± 2 min) | 7:00 PM<br>(63 ± 27 min) | 4:00 PM<br>(51 ± 10 min) |

Abbreviations: RPE, rating of perceived exertion; GPS, global positioning system

#### Paragraph 9

Throughout the study (i.e., 9 consecutive days of the tournament), the subjects were hosted in the same hotel. The players slept in twin rooms with separate beds. All meals were eaten at the same hotel restaurant (i.e., breakfast, lunch, and dinner). Similarly, all training sessions were conducted within the hotel’s sports complex. All training sessions and matches were performed on an outdoor natural grass pitch. Ambient temperature ranged from 16–18°C during the day and 10–12°C during the night. The competitive matches were held in 3 different stadiums located in the same district (Faro, Portugal; the furthest stadium from the hotel was ∼1h by bus). Therefore, long journeys and the consequences (e.g., travel fatigue and jet leg) were avoided. Training schedules were set at the discretion of the team coaching staff. There was no interference by the research team in the athletes’ regular training schedule or sleep/wake patterns. The athletes were free to consume snacks, nutritional supplements, and caffeine during the data collection period.

#### Paragraph 10

Technical problems and/or player compliance resulted in some missing data points (sleep variables: valid cases, n=142 [92%], and missing cases, n=13 [8%]; nocturnal HRV indices: valid cases, n=137 [88%], and missing cases, n=18 [12%]; TD, exposure time and HSR: valid cases, n=137 [88%], and missing cases, n=18 [12%]; s-RPE: valid cases, n=103 [67%], and missing cases, n=52 [34%]).

### Measurements

#### Paragraph 11. Training and match load monitoring

Individual training and match exposure time was routinely recorded by a member of the team’s medical staff. Subsequently, the recorded exposure time was used to select the actual global positioning system (GPS) training data. The same procedure was used during matches, based on the individual effective playing time, including substitute players. The amount of activities performed during the tournament was monitored using wearable 18-Hz GPS units (STATSports Apex, Northern Ireland). The accuracy of this device has been previously examined, reporting a nearly perfect criterion validity to measure distance during team sport specific movements (ICC = 0.98)[39]. A special vest was tightly fitted to each player, with the receiver placed between the scapulae. All devices were activated 15-min before the data collection to allow acquisition of satellite signals in accordance with the manufacturer’s instructions. In addition, in order to avoid inter-unit error, each player wore the same GPS unit throughout the tournament [40]. Data were subsequently downloaded and adjusted to training and match exposure using corporate software (STATSports Apex, Northern Ireland). TD covered was adopted as the measure of training and match volume. High-speed running (HSR, >12.6 km·h^-1^) was adopted as the measure of high-intensity activity performed, according to a recent study in top-level female players [41].

#### Paragraph 12

Psychophysiological response to exercise was quantified using the session-rating of perceived exertion method (s-RPE). Throughout the tournament, the players reported individual RPE using the Borg category ratio scale (CR10) via a customized mobile application, approximately 30 min after each training session or match. The CR10 score (perceived intensity) was subsequently multiplied by individual exposure time (training and match volume), thus providing an overall load quantification of the session or match [42].

#### Paragraph 13. Sleep monitoring

Night-sleep was assessed using 3-axial accelerometers (Actigraph LLC wGT3X-BT, Pensacola, USA) worn on the non-dominant wrist. Wrist-worn accelerometers have been used to monitor sleep in elite athletes [15], and validated against polysomnography (PSG; [43]). Data were analysed using corporate software (ActiLife LLC Pro software v6.13.3, Pensacola, USA). The sampling frequency was 50 Hz and the epoch of activity counts was 60 s [44]. All sleep variables were determined every night throughout the tournament using the Sadeh’s algorithm [44]. Objective sleep measures included total sleep time (total amount of sleep obtained), time in bed (time between lying down until getting up the next day), wake-up time (time between the last minute of sleep and getting up from bed), sleep onset time (time of the first epoch of sleep between time of trying to initiate sleep and time at wake up), wake after sleep onset (number of min awake after sleep onset), sleep fragmentation index (sum of mobility and immobility accesses in one minute, divided by the number of immobility accesses), latency (time in minutes attempting to fall asleep), and sleep efficiency (percentage of time in bed that was spent asleep) [44]. It is important to mention that although some studies have reported SE ≥85% as insufficient sleep quality [4], according to the latest National Sleep Foundation report [16], SE ranging from 75-84% is considered uncertain for young adults/adults, whereas SE ≤74% indicates inappropriate sleep quality for the respective age category. Therefore, sleep quantity (TST) and sleep quality (SE) were analysed according to the Sleep National Foundation report (i.e., TST <7h; 420 min as an indicator of inappropriate sleep quantity, and SE ≤74% as inappropriate sleep quality) [16].

#### Paragraph 14. Cardiac autonomic activity monitoring

The slow-wave sleep episode (SWSE) method, which accounts for the deep stage of sleep [32], was used to analyse the cardiac autonomic activity during night-sleep [45]. This method records 10 minutes of normal RR intervals, considering the criteria proposed by Brandenberger et al., (2005). HR monitors (Firstbeat Bodyguard2^®^, Firstbeat Technologies, Finland) were used during sleep. This device has been validated against standard electrocardiogram equipment to detect heartbeats [46]. The RR intervals analyzed in time domain measures included mean RR interval (mRR), mean HR, RMSSD (square root of the mean of the sum of the squares of differences between adjacent normal RR intervals; vagal modulation index), and SDNN (standard deviation of all NN [RRintervals] interval). RR intervals were also used to produce the Poincaré plot SD_1_ (short-term beat-to-beat variability) and SD_2_ (long-term beat-to-beat variability) values. Fast Fourier Transform (Welch’s periodogram: 300-s window with 50% overlap)[47] was used to obtain measures of nocturnal cardiac autonomic activity in the frequency domain, considering both LF (0.004–0.15 Hz) and HF (0.15–0.4 Hz) indices. The ratio (i.e., LF/HF) index was calculated from the non-transformed LF and HF data [47]. In addition, to reduce any potential non-uniformity or skewness in HRV, data were log-transformed by taking the natural logarithm (ln) before conducting any statistical analyses [48]. In all cases, HRV was calculated using Kubios HRV 3.0.0® software (Kubios Oy, Kuopio, Finland).

#### Paragraph 15

Buchheit [32] proposed 3% as the fixed smallest worthwhile change (SWC) to detect eventual changes related to positive and negative adaptations in HRV-derived indices. Given that both 0.5×CV [32] and 1×CV have been acceptably used to account for within-athlete variations in HRV-derived indices, it is possible to state that using 3% as proposed by Buchheit [32] is acceptable and conservative for lnRMSSD, especially for soccer players [26].

#### Paragraph 16. Statistical Analysis

Sample distribution was tested using the Shapiro–Wilk test for sleep patterns, nocturnal cardiac autonomic activity indices (HRV), and training and match load variables for each day of the tournament. Sleep patterns, nocturnal autonomic activity, and training load variables displayed during the 9 days of the tournament are presented as mean ± SD for the indices that displayed normal distribution and presented as median (interquartile range) for data that did not present normal distribution. The coefficient of variation (CV; CV=([SD/mean] ×100) was calculated for the whole group and also intra-individually for TST and lnRMSSD indices across 9 days of the international tournament. The SWC was calculated from the intra-individual CV of lnRMSSD (lnRMSSD_CV_), considering the 9 days of the tournament [32].

#### Paragraph 17

A within-subjects linear mixed model analysis was performed to examine differences in TST, SE, and nocturnal lnRMSSD variables across 9 days of the tournament. An α-level of 0.05 was set as the level of significance for statistical comparisons. The days of the tournament (i.e., DT, DM, and EM) were included as a fixed effect and player identity (subject ID) as the random effect. Furthermore, among the recommended variance-covariance structure models, compound symmetry was selected according to the smallest Akaike Information Criterion assessment [49], based on the Restricted Maximum Likelihood method. Pairwise comparisons (Bonferroni) were used to show the day-to-day mean differences for TST, SE, and nocturnal lnRMSSD indices.

## Results

### Paragraph 18. Training and match load variables

As a group, the s-RPE, TD, exposure time, and HSR during training sessions ranged from 131 to 360 arbitrary units (AU), 2201 to 4284 m, 34 to 76 min, and 130 to 756 m, respectively. During matches these variables ranged from 504 to 602 AU, 7012 to 7746 m, 64 to 83 min, and 1678 to 1888 m, respectively. These data are summarized in Figure1 (Fig 1. A, B, C, and D).

**Figure 1.**
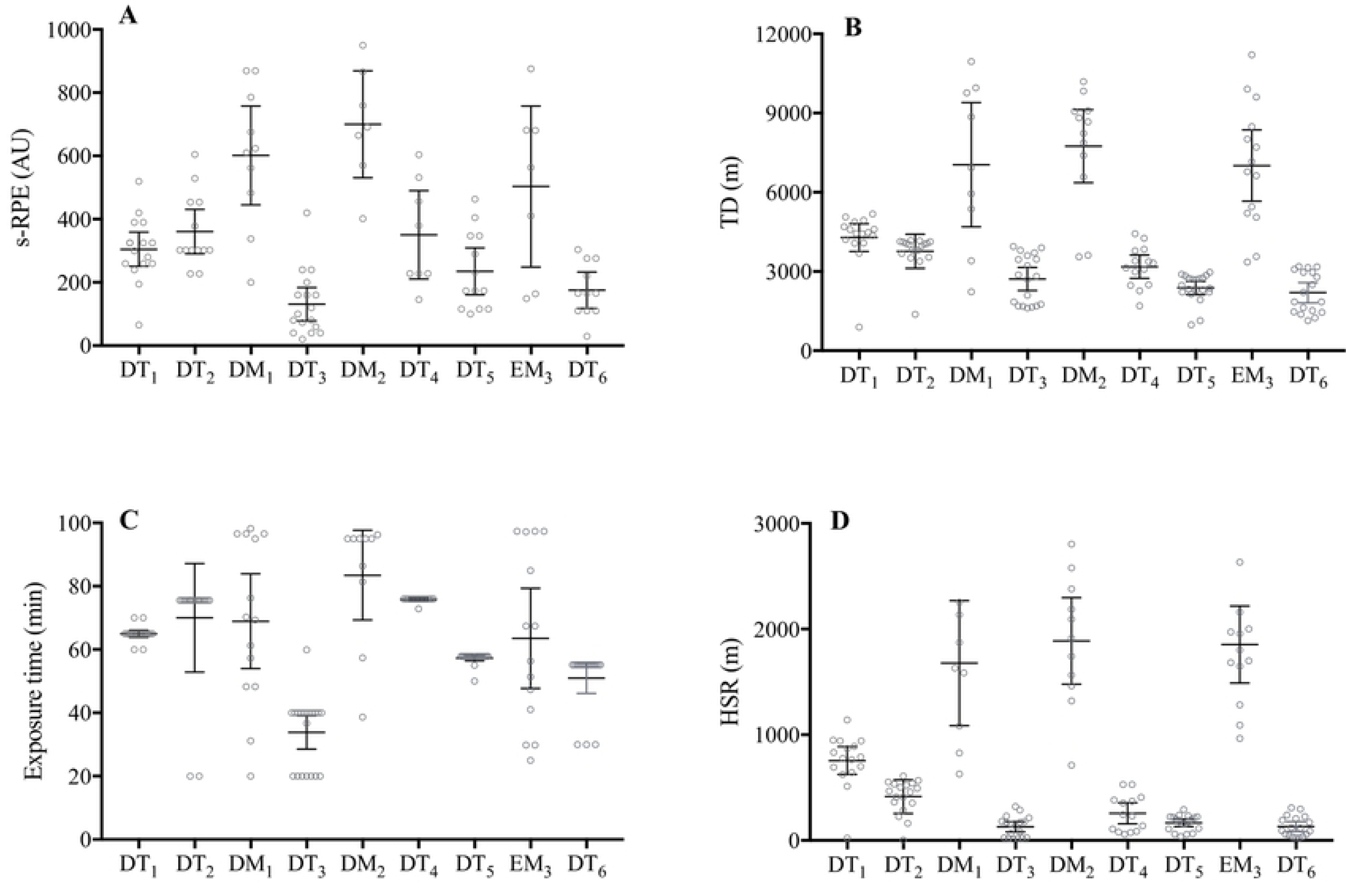
Individual subject session of perceived exertion (A), total distance (B), exposure time (C), and high-speed running (D) responses during 9 consecutive days of an international tournament in elite female soccer players. Group mean ± 95% confidence interval are presented (black lines) for s-RPE and HSR, and median (interquartile range) for TD and exposure time (black lines) for each day in elite female soccer players. Abbreviations: s-RPE, session-rating of perceived exertion; TD, total distance; HSR, high-speed running; DT, day-training; DM, day-match; EM, evening-match. Note: s-RPE: DT_1_ (n=15); DT_2_ (n=14); DM_1_ (n=10); DT_3_ (n=17); DM_2_ (n=7); DT_4_ (n=8); DT_5_ (n=13); EM_3_ (n =7); and DT_6_ (n=11). TD, exposure time, and HSR: DT_1_ (n=16); DT_2_ (n=18); DM_1_ (n=9); DT_3_ (n=18); DM_2_ (n=11); DT_4_ (n=14); DT_5_ (n=20); EM_3_ (n=13); and DT_6_ (n=18).

### Paragraph 19

Individually, the s-RPE, TD, exposure time, and HSR during training sessions ranged from 20 to 680 AU, 892 to 5176 m, 20 to 76 min, and 80 to 1140 m, respectively. During matches these values ranged from 149 to 876 AU, 2236 to 11210 m, 20 to 98 min, and 629 to 3213 m, respectively. These data are summarized in Figure1 (Fig 1. A, B, C, and D).

### Paragraph 20. Sleep variables

As a group, player TST ranged between 7:41±0:48 and 8:26±0:41 hours during all DT and both DM of the tournament, Table 2. However, a lower duration of TST was observed after EM (6:47±0:58 hours) compared to all days of the tournament (*p*<0.001), especially when compared to both DM_1_ and DM_2_ (−1:39±0:17 hours and −1:28±0:12 hours, respectively; *p*<0.001). In addition, a lower SE was found after EM compared to all days of the tournament (*p*<0.001), especially when compared to both DM_1_ and DM_2_ (−6±3% and −4±2%, respectively; *p*<0.001). All the sleep-related variables are presented in Table 2.

**Table 2.** Actigraphy sleep responses during 9 days comprising an international tournament in elite female soccer players. Values are mean ± standard deviation (SD) and median (interquartile range).

|  | DT<sub>1</sub> (n=20) | DT<sub>2</sub> (n=20) | DM<sub>1</sub> (n=13)<br>missing<br>(n=1; 7%) | DT<sub>3</sub> (n=19)<br>missing<br>(n=1; 5%) | DM<sub>2</sub> (n=13)<br>missing<br>(n=1; 7%) | DT<sub>4</sub> (n=12)<br>missing<br>(n=2; 14%) | DT<sub>5</sub> (n=17)<br>missing<br>(n=3; 15%) | EM<sub>3</sub> (n=13)<br>missing<br>(n=2; 13%) | DT<sub>6</sub> (n=15)<br>missing<br>(n=3; 16%) |
| --- | --- | --- | --- | --- | --- | --- | --- | --- | --- |
| Sleep onset time (h:min) | 23:31±0:23* | 23:12 ± 0:36* | 23:14 ± 0:34* | 23:26 ± 0:34* | 23:19 ± 0:56* | 23:48 ± 0:39* | 23:30 ± 0:33* | 0:45 ± 0:32 | 23:35 ± 0:39* |
| Wake up time (h:min) | 8:27 ± 0:31 | 8:39 (0:36) | 9:02 (1:01) | 8:38 (0:37) | 8:25 ± 0:22 | 8:14 ± 0:27 | 8:37 ± 0:27 | 8:56 ± 0:34 | 8:39 (0:43) |
| TIB (h:min) | 9:06 ± 0:45 | 9:15 (0:58) | 9:42 ± 0:28 | 9:03 ± 0:32 | 9:08 ± 1:11 | 8:21 ± 0:44 | 8:54 ± 0:56 | 7:50 ± 0:46 | 8:34 ± 0:33 |
| TST (h:min) | 7:56 ± 0:47* | 7:58 ± 0:54* | 8:26 ± 0:41* | 7:54 ± 0:36* | 8:15 ± 1:08* | 7:41 ± 0:48* | 8:02 ± 0:57* | 6:47 ± 0:58 | 7:55 ± 0:38* |
| WASO(min) | 44 (32) | 59 (38) | 61 ± 32 | 45 (37) | 41 (35) | 42 ± 25 | 53 ± 33 | 65 ± 35 | 63 (74) |
| SFI (%) | 26 ± 9 | 26 ± 9 | 26 ± 8 | 29 ± 11 | 27 ± 10 | 23 ± 10 | 27 ± 9 | 32 ± 10 | 24 (18) |
| SE (%) | 91 ± 6* | 91 ± 6* | 87 ± 4* | 91 ± 6* | 85 ± 5* | 90 ± 6* | 92 ± 6* | 81 ± 7 | 91 ± 4* |
| Sleep latency(min) | 4 ± 3 | 3 (2) | 7 ± 2* | 3 (2) | 9 ± 4* | 4 ± 3 | 3 (2) | 20 ± 9 | 4 ± 3 |

\*Significantly different compared to EM (P<0.001)
Abbreviations: DT, day-training; DM, day-match; EM, evening-match; TIB, time in bed; TST, total sleep time; WASO, wake after sleep onset;
SFI, sleep fragmentation index; SE, sleep efficiency.
Note: The mean values represent only the players that trained and played on the respective day of the tournament (goalkeepers were excluded).

### Paragraph 21

Individually, some players slept less than recommended (<7 hours; 420 min) after DT_1_ (n=4; player 8,13,19 and 20), DT_2_ (n=3; player 6, 13 and 20), DM_1_ n=0, DT_3_ (n=1; player 12), DM_2_ (n=6; 8, 10, 16 and 20), DT_4_ (n=2; player 12 and 13), DT_5_ (n=2; player 2 and 12), EM_3_ (n=8; player 5, 6, 8, 10, 12, 16, 17 and 20), and DT_6_ (n=1; player 20); especially after EM (TST range between 6:00-6:54 h). TST_CV_ ranged between 3.1 and 18.7 % (Figure 2). Overall, all players presented good sleep quality (i.e., sleep efficiency ≥75%; individual range between 75-98%) across all days of the tournament (Figure 2).

**Figure 2.**
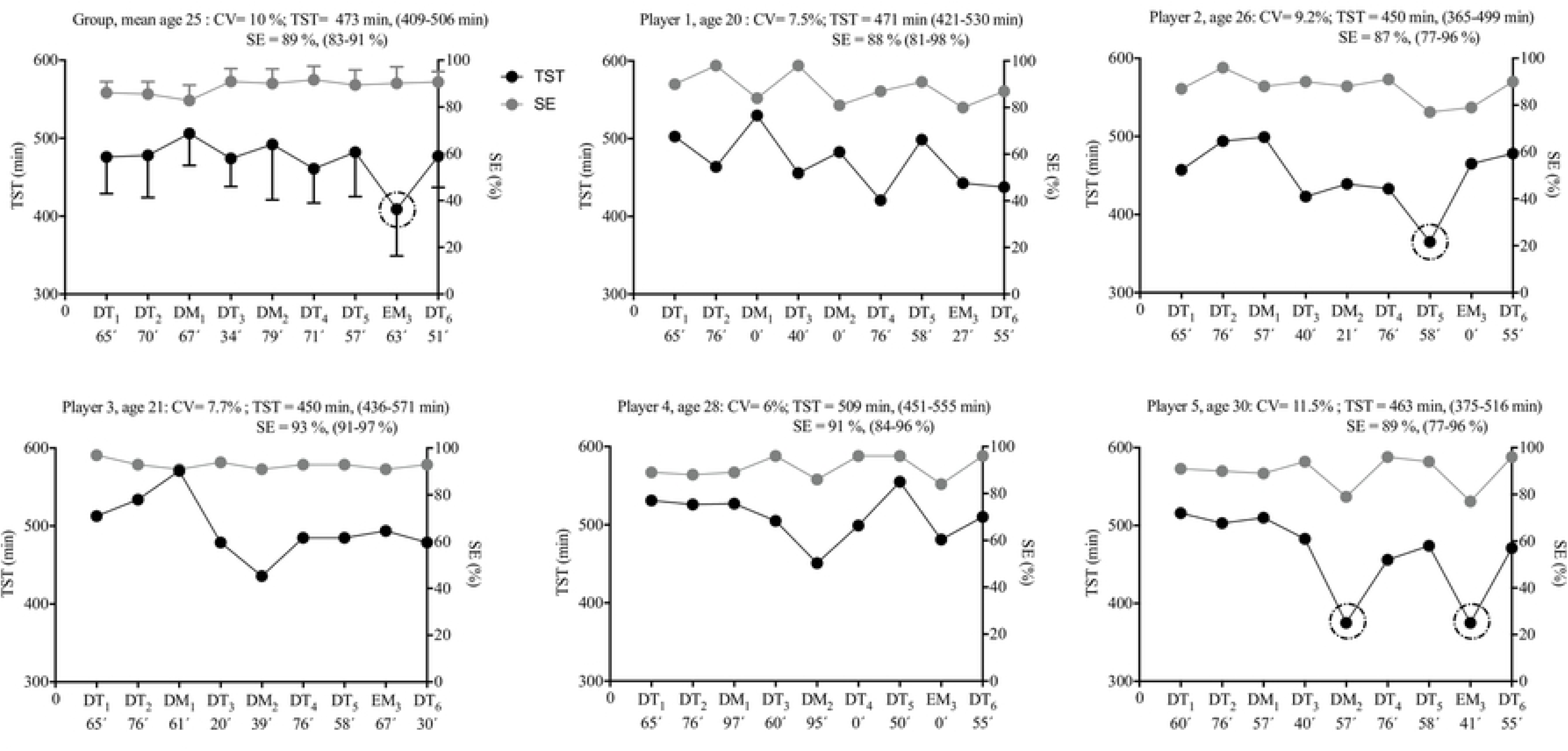
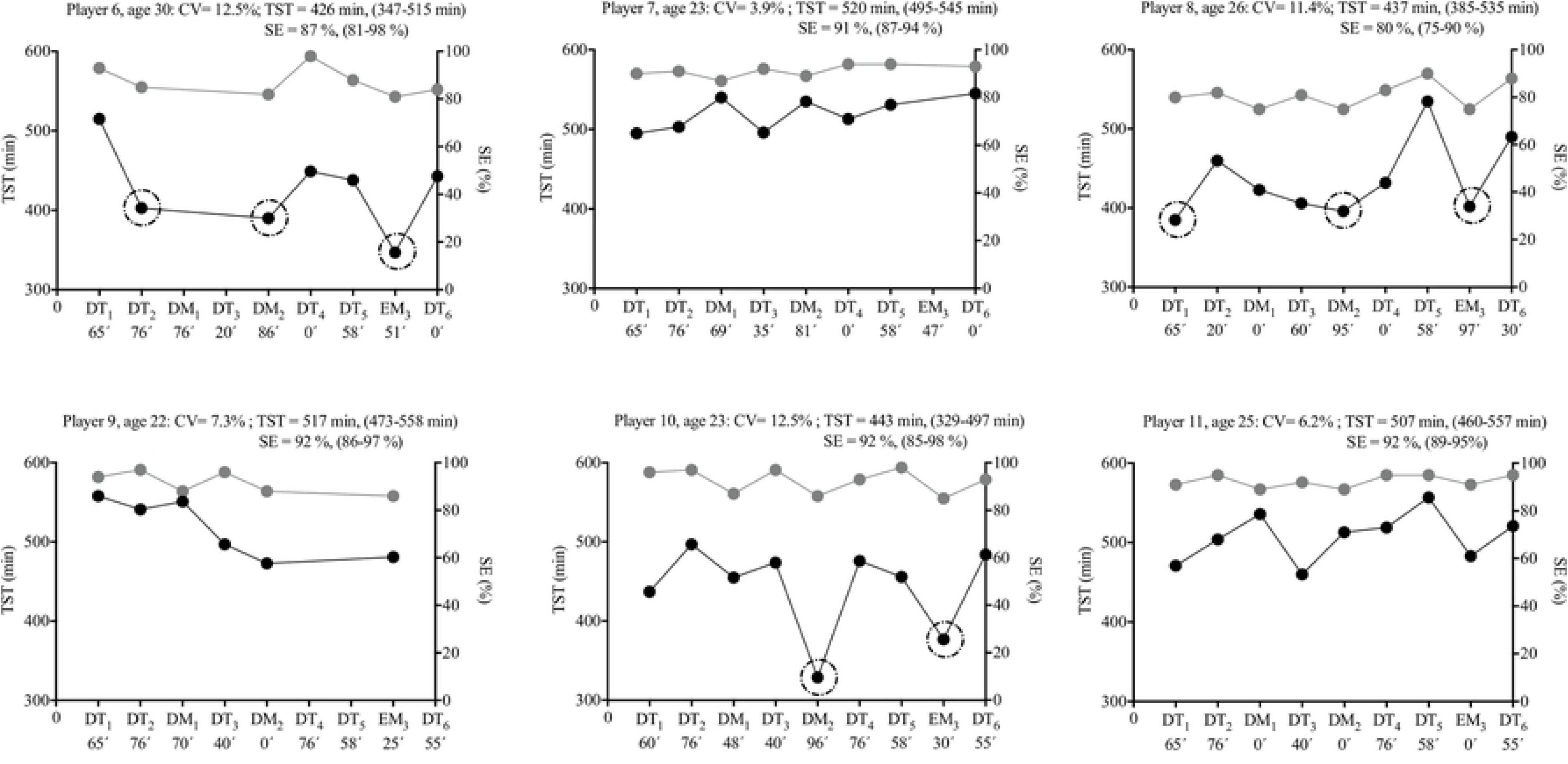
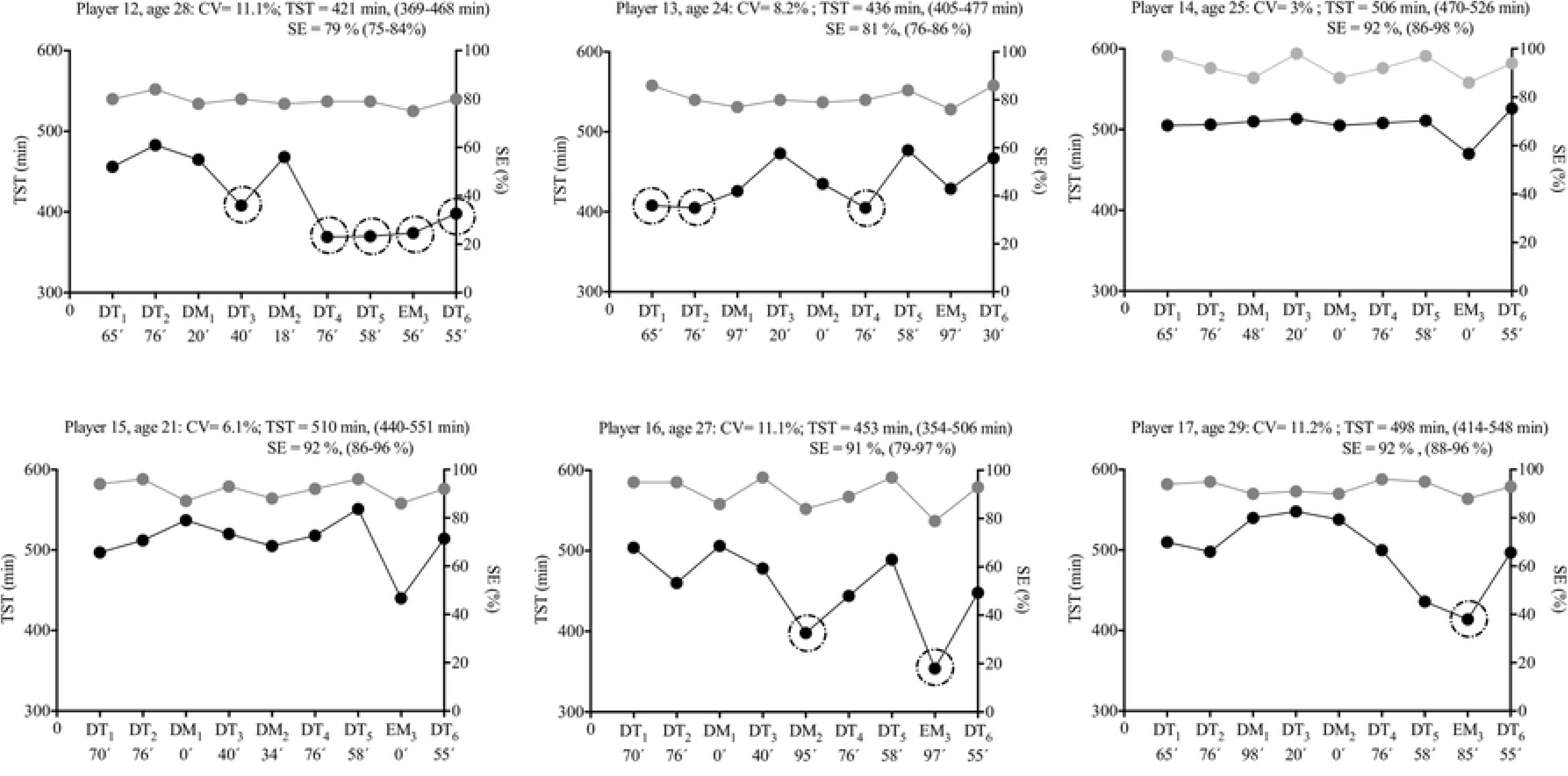
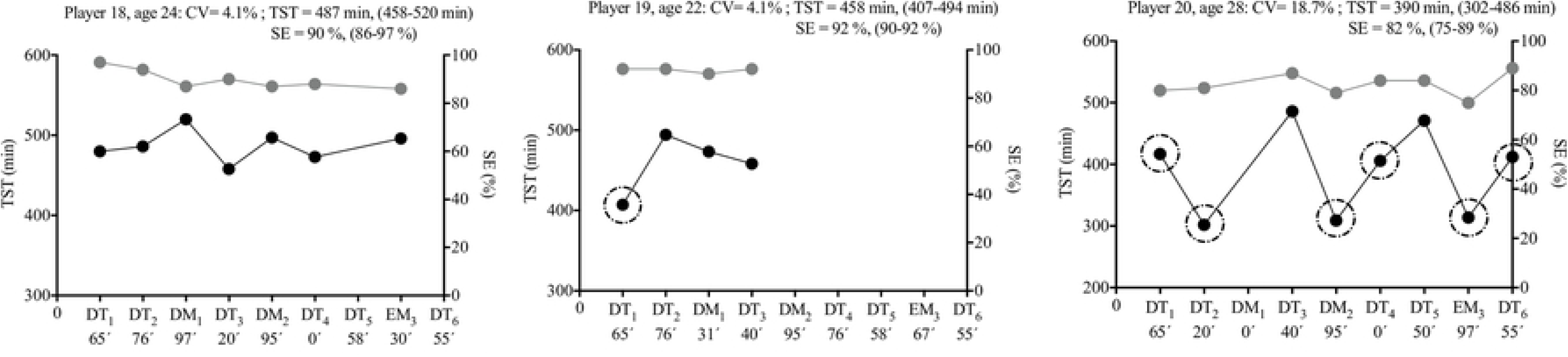
Group and individual subject (n=20) total sleep time (TST) and sleep efficiency (SE) displayed during 9 days of an international tournament. Group and individual black and grey dots represent daily changes in TST and SE, respectively. TST and SE averages (with max-min values) are also presented. The black dashed circumferences represent the days that TST was lower than recommended (i.e., <420 min; <7 hours). The exposure time (min) for each day is also presented on all days of the tournament (x axis). Abbreviations: DT, day-training; DM, day-match; EM, evening-match; CV, coefficient of variation.

### Paragraph 22. Cardiac autonomic activity variables

As a group, nocturnal autonomic cardiac activity was not affected during the 9 days of the tournament. Overnight lnRMSSD ranged from 4.19±0.88 to 4.54±0.42 ln[ms] (*p*>0.05). Nocturnal autonomic cardiac activity data are presented in Table 3.

**Table 3.** Overnight cardiac autonomic activity (SWSE method) responses during 9 days of the international tournament in elite female soccer players. Values are mean ± standard deviation (SD) and median (interquartile range).

|  | DT<sub>1</sub> (n=19)<br>missing<br>(n=1; 5%) | DT<sub>2</sub> (n=18)<br>missing<br>(n=2; 10%) | DM<sub>1</sub> (n=13)<br>missing<br>(n=1; 7%) | DT<sub>3</sub> (n=19)<br>missing<br>(n=1; 5%) | DM<sub>2</sub> (n=12)<br>missing<br>(n=2; 14%) | DT<sub>4</sub> (n=12)<br>missing<br>(n=2; 14%) | DT<sub>5</sub> (n=16)<br>missing<br>(n=4; 20%) | EM<sub>3</sub> (n=12)<br>missing<br>(n=3; 20%) | DT<sub>6</sub> (n=15)<br>missing<br>(n=3; 17%) |
| --- | --- | --- | --- | --- | --- | --- | --- | --- | --- |
| Heart rate (bpm) | 46.5 ± 5.2 | 47.4 ± 7.0 | 46.1 ± 7.3 | 46.9 ± 6.9 | 48.3 ± 4.7 | 46.0 ± 6.5 | 46.9 ± 6.0 | 48.8 ± 5.6 | 47.5 ± 7.1 |
| RR interval (ln[ms]) | 7.2 ± 0.1 | 7.2 ± 0.2 | 7.2 ± 0.2 | 7.2 ± 0.1 | 7.1 ± 0.1 | 7.2 ± 0.1 | 7.2 ± 0.1 | 7.1 ± 0.1 | 7.2 ± 0.2 |
| RMSSD (ln[ms]) | 4.4 ± 0.7 | 4.2 ± 0.9 | 4.4 ± 0.7 | 4.3 ± 0.7 | 4.4 ± 0.6 | 4.5 ± 0.4 | 4.3 ± 0.5 | 4.4 ± 0.7 | 4.5 ± 0.4 |
| SDNN (ln[ms]) | 4.0 ± 0.7 | 3.9 ± 0.8 | 4.0 ± 0.6 | 3.9 ± 0.7 | 4.1 ± 0.6 | 4.2 ± 0.3 | 4.0 ± 0.4 | 4.1 ± 0.6 | 4.2 ± 0.3 |
| SD1 (ln[ms]) | 4.0 ± 0.7 | 3.8 ± 0.9 | 4.0 ± 0.7 | 4.0 ± 0.7 | 4.1 ± 0.6 | 4.2 ± 0.4 | 4.0 ± 0.5 | 4.1 ± 0.7 | 4.2 ± 0.4 |
| SD2 (ln[ms]) | 3.9 ± 0.6 | 3.8 ± 0.7 | 4.0 ± 0.5 | 3.9 ± 0.9 | 4.0 ± 0.5 | 4.1 ± 0.3 | 3.9 ± 0.3 | 4.0 ± 0.5 | 4.1 ± 0.3 |
| LF (ln[ms<sup>2</sup>]) | 6.5 ± 1.2 | 6.3 ± 1.4 | 6.9 ± 1.1 | 6.5 ± 1.5 | 6.8 ± 0.9 | 7.0 ± 0.6 | 6.4 ± 0.6 | 6.9 ± 1.0 | 6.9 ± 0.7 |
| HF (ln[ms<sup>2</sup>]) | 7.4 ± 1.4 | 7.1 ± 1.7 | 7.6 ± 1.5 | 7.3 ± 1.5 | 7.5 ± 1.2 | 7.7 ± 0.8 | 7.3 ± 0.9 | 7.5 ± 1.4 | 7.7 ± 0.8 |
| LF/HF | 0.7 (0.5) | 0.6 (0.5) | 0.8 (0.5) | 0.6 (0.5) | 0.6 ± 0.4 | 0.6 (0.5) | 0.5 (0.4) | 0.7 ± 0.5 | 0.4 (0.2) |

Abbreviations: DT, day-training; DM, day-match; EM, evening-match; RR interval, variations between consecutive heart beats (beat to beat); RMSSD, square root of the mean of the sum of the squares of differences between adjacent NN intervals; SDNN, standard deviation of all NN interval; SD1, short-term beat-to beat variability; SD2, long-term beat-to-beat variability; LF, low frequency; HF, high frequency; LF/HF, ratio of the low-to high frequency power.
Note: The mean values represent only the players that trained and played on the respective day of the tournament (goalkeepers were excluded).

### Paragraph 23

Individually, two players (players 6 and 7) appeared to present higher lnRMSSD_CV_ (11.7 and 11.5%; respectively), which occurred simultaneously with a reduced lnRMSSD average (3.66 and 2.73 ms; respectively), throughout the tournament, in contrast to the remaining team (individual lnRMSSD ranging between 3.91 and 5.37 ms, and lnRMSSD_CV_ ranging between 2.8 and 9%) (Figure 3).

**Figure 3.**
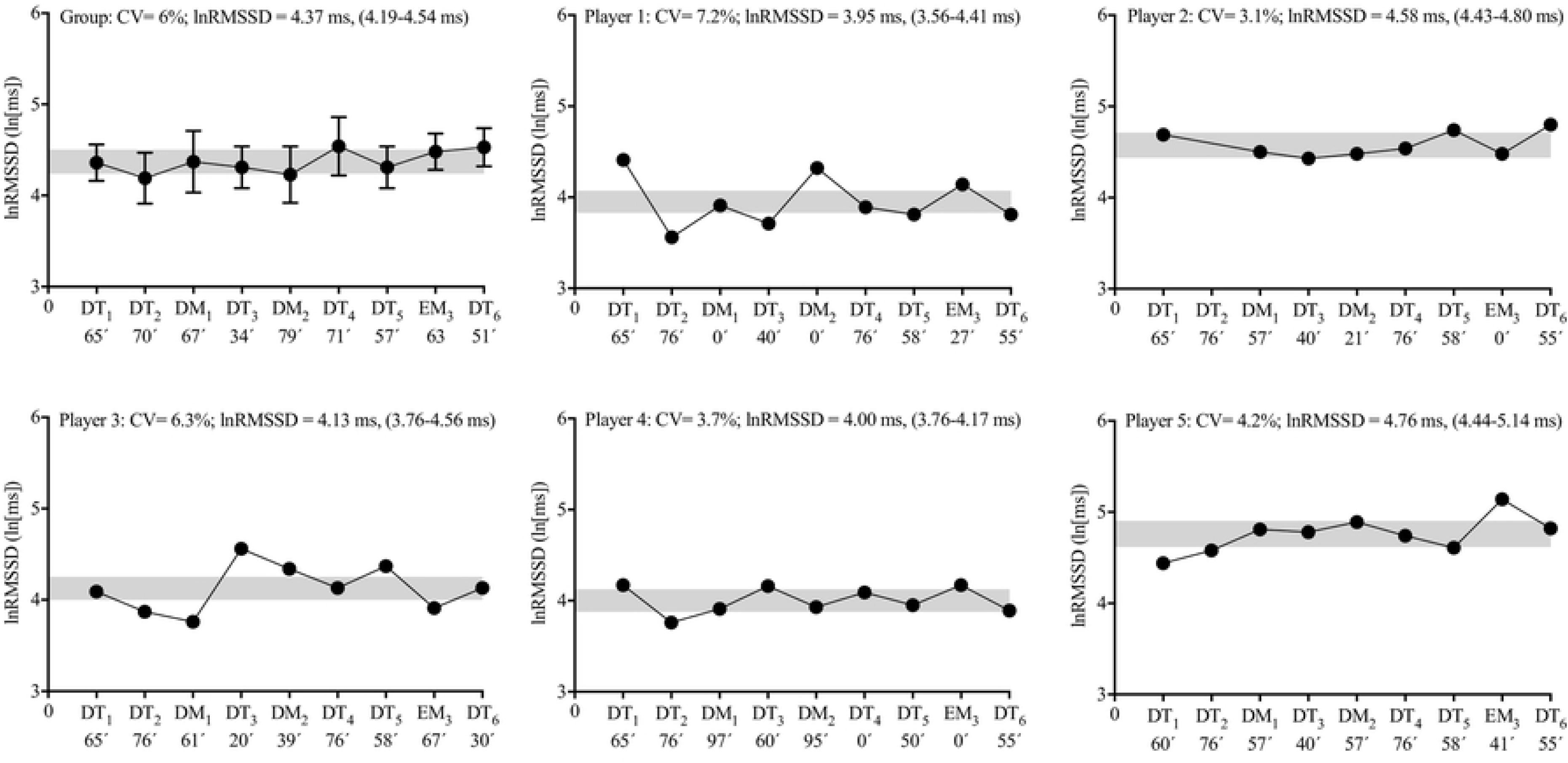
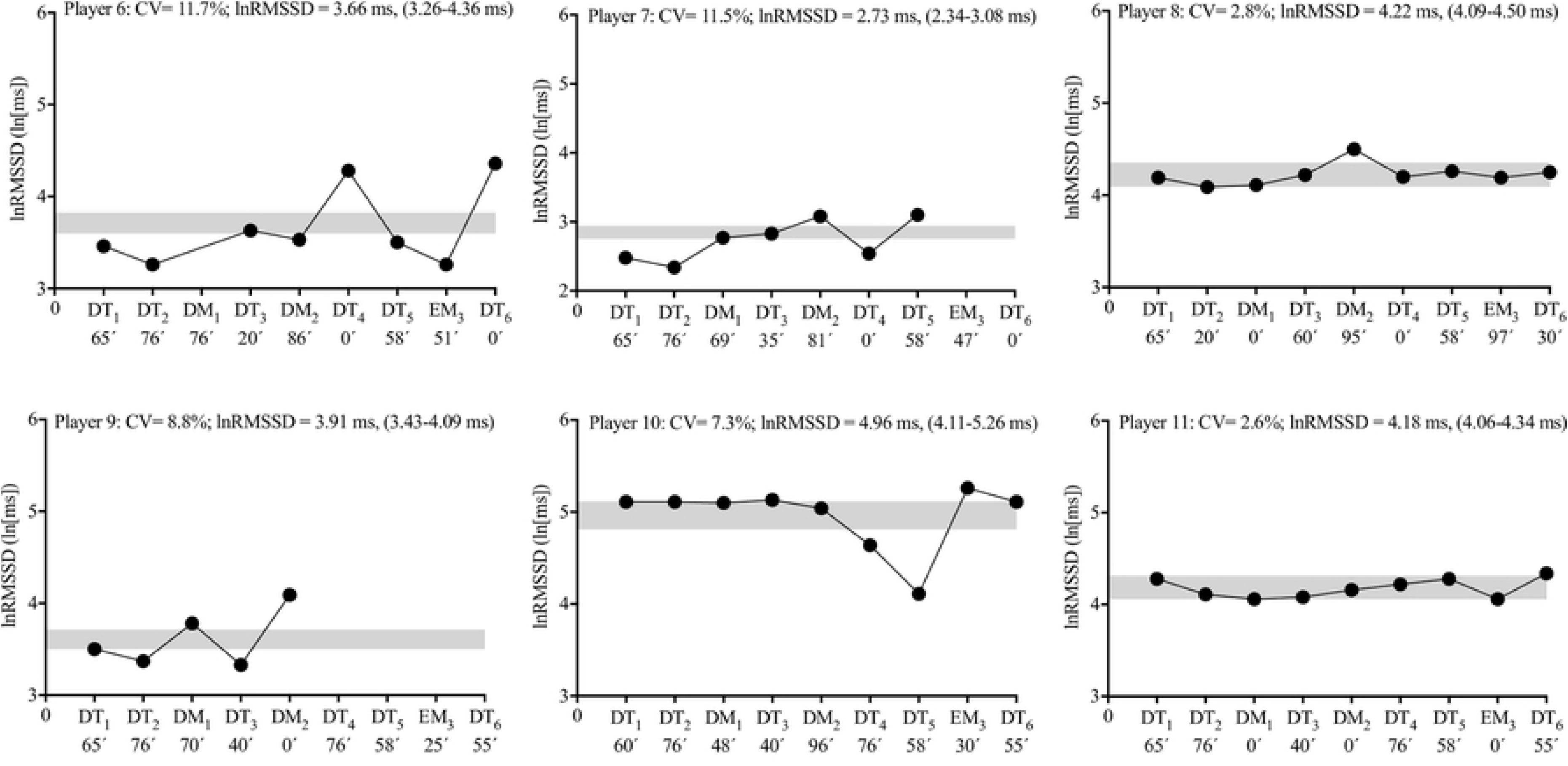
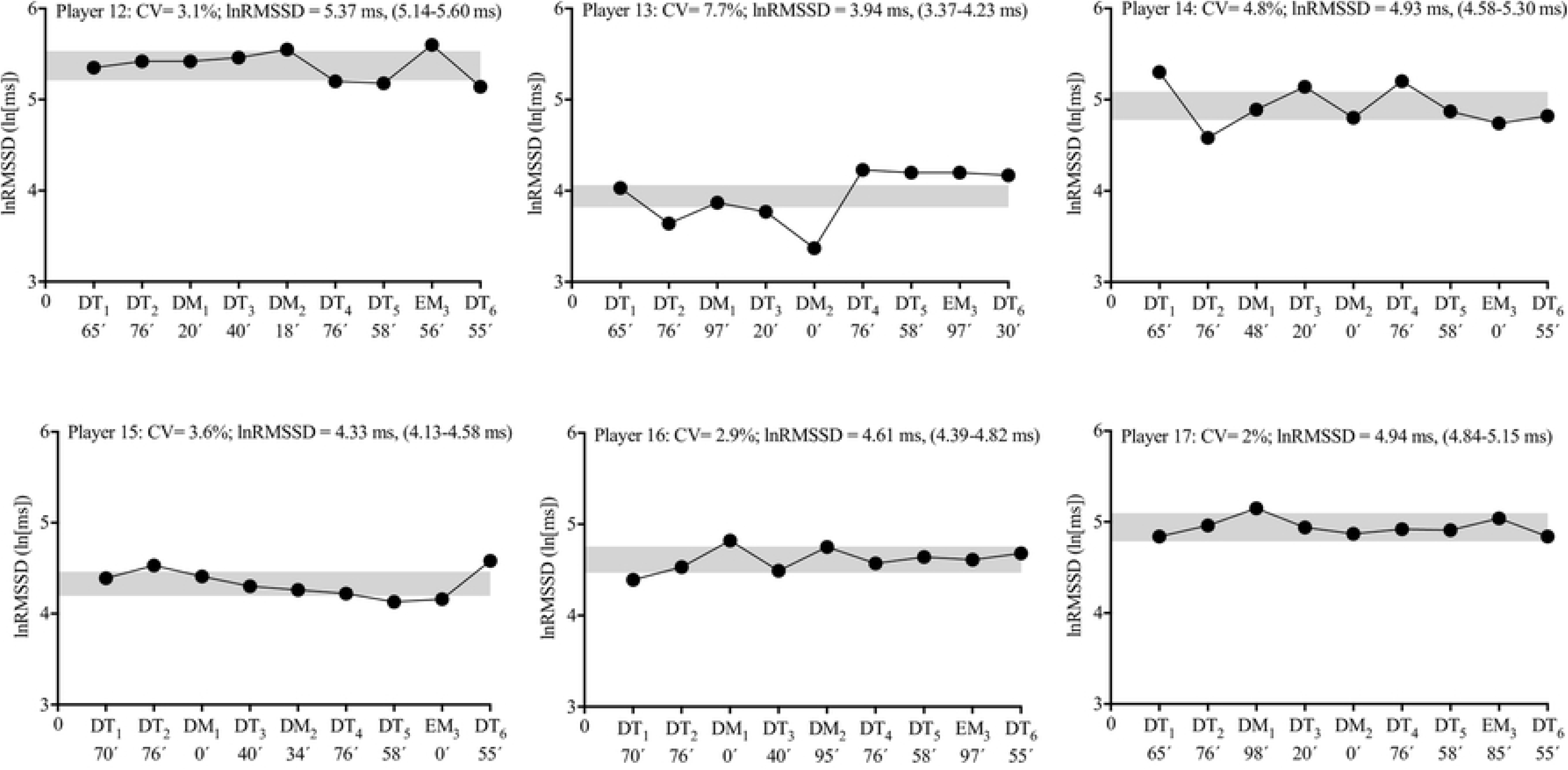
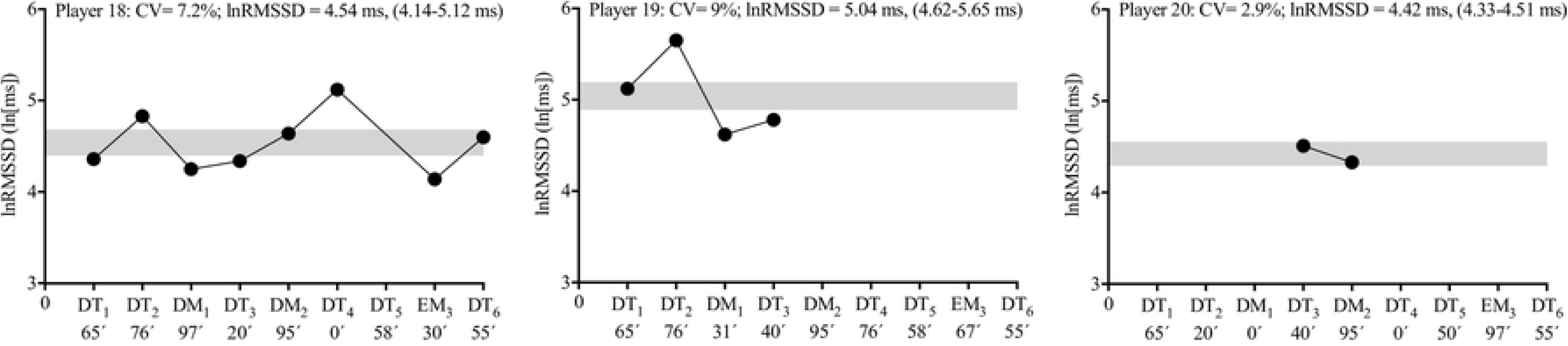
Group and individual subject (n=20) cardiac parasympathetic activity displayed during 9 days of an international tournament. Group and individual black dots represent daily changes in natural logarithm of the root mean square of successive RR intervals (lnRMSSD). lnRMSSD averages (with max-min values) are also presented. The grey area represents the smallest worthwhile change zone (3%) [32]. The exposure time (min) for each day is also presented on all days of the tournament (x axis). Abbreviations: DT, day-training; DM, day-match; EM, evening-match; CV, coefficient of variation.

## Discussion

### Paragraph 24

The main finding from the sleep results was that some players presented less TST than recommended during different days of the tournament (i.e., independently of being DT, DM and EM). However, after EM, a higher number of athletes slept for less than 7 hours in contrast to the remaining days of the tournament during which they presented adequate sleep duration. Nevertheless, all players displayed good sleep quality.

### Paragraph 25

The main finding from overnight HRV results was that most players seemed to present small fluctuations in nocturnal cardiac autonomic activity, while two players appeared to present higher lnRMSSD_CV,_ occurring simultaneously with a reduced lnRMSSD average during the tournament, compared to the remaining players of the team.

### Paragraph 26

Overall, as a group, players accumulated adequate sleep quantity and presented good sleep quality on all training days and DM of the tournament. However, significantly decreased durations of TST and SE were observed after EM compared to DM of the tournament. These results are in accordance with recent studies showing that soccer players presented sleep duration within the appropriate healthy range on training days and match days concluded before 6 PM, but slept significantly less, delayed bedtime, and presented a lower SE after matches starting after 6 PM [50, 51]. In fact, the impact of night matches (i.e., schedule time) on subsequent sleep is well established [52, 53]. For example, Sargent and Roach (2016) examined the sleep of elite Australian football players on the night immediately following a day match or following an evening match. During the night immediately after the evening match, the players initiated sleep 2.5 hours later and obtained 2.1 hours less sleep when compared to sleep immediately following the day match. In the present study, following EM, female soccer players initiated sleep much later compared to DM_1_ and DM_2_, and obtained less sleep quantity compared to DM_1_ and DM_2_. Another interesting result was that higher sleep latency was found after EM compared to DM_1_ and DM_2_. These results are in accordance with a large epidemiological survey that reported that 24% of respondents perceived difficulty in initiating sleep after late-evening exercise compared to 7% after early evening exercise [54]. Recently, it was also shown that sleep latency in high-level female soccer players was negatively affected after night-training sessions (start 9:00 PM) compared to resting days and match days (3:00 PM) [18]. These results suggest that late-evening exercise, occurring close to bedtime sleep, can impose a high risk of obstructing falling asleep at night. Thus, an acceptable explanation for the results observed in the present study could be the time scheduling of the matches, suggesting the potential value for sleep education strategies and interventions to promote appropriate sleep and recovery, especially after night competitions, since athletes are often unable to achieve recommended TST and SE, with lower values on the night of competition compared with the previous night [4].

### Paragraph 27

A main finding from the individual sleep analysis was that some players slept less than recommended during the tournament, especially after EM. Importantly, seven out of eight players who played the EM also presented shorter TST on the night after EM (start 7 PM) compared with the previous night (DT). As already mentioned, these results are in accordance with a recent study [4] in elite athletes that showed nocturnal TST was shorter on the night after the match (start ≥6 PM) compared with the previous night (training and match or rest day).

### Paragraph 28

Besides the training schedule, another possible explanation for the sleep results could be the training and match loads that players were exposed to during the tournament, especially during matches [4]. In fact, on days DM_2_ and EM, more players that participated during the match slept less than recommended, with both days presenting the highest values of external load (e.g., HSR) and exposure time during the tournament. Interestingly, no sleep disturbances were found for any players that played DM_1_. Nonetheless, despite the external training and match loads that players were exposed to, each player seemed to present good sleep quality across all days of the tournament. These results could also be supported by the appropriate values of sleep latency observed for all players during the tournament. Additionally, in the current study, the within-player variability in TST (CV=7.4%) was relatively low compared with a recent study that found a good within-player consistency of sleep (CV=15.2%), across 8 days of rest (without exercise) and 8 days of night-training sessions in highly trained female soccer players [19]. Thus, even under stress imposed by tournament scheduling and training and match loads, the players maintained relatively good consistency in sleep habits to recover from the training sessions and matches.

### Paragraph 29

Notably, six players seemed to have constantly “compensated” the low TST after participating in matches (independently of being DM or EM), including one player that played all matches (player 10), by extending sleep duration on the next day, probably as a “self-sleep hygiene” strategy. These findings are relatively consistent with previous data suggesting that training and competition schedules dictate the sleep/wake behavior of elite athletes [7, 9, 55]. Unfortunately, the strategies that athletes used to increase their sleep duration during the day (e.g., naps) were not recorded which is recognized as a limitation of the current study. Therefore, the findings from the present study indicate that, if given the opportunity, elite female soccer players will compensate by extending their sleep the following day to ameliorate any sleep loss and fatigue [11]. Taken together, these findings are of considerable importance for athletes and coaches because nights of reduced sleep (i.e., 5–7 h of sleep per night) can lead to deficits in neurobehavioural performance and increases in subjective feelings of sleepiness and fatigue [56]. Therefore, if the training or competition schedule provides sufficient space for recovery strategies, extending bedtime to achieve prolonged sleep duration appears to be a beneficial approach. As prevalence of sleep restriction among athletes seems to be high [57], athletes are encouraged to implement 30 to 60 minutes of additional sleep each night as a ‘self-experiment’ which should be monitored and supported by staff members [58]. In addition, when feasible, athletes may also achieve additional sleep time by implementing daytime naps. Furthermore, habitual napping can be generally encouraged, whereas timing (i.e., preferably in the early afternoon) and duration (i.e., < 30 min) should be appropriately planned [59]. However, naps shorter than one hour are recommended, and not too close to bedtime as it may interfere with sleep [59].

### Paragraph 30

In the current study, no significant changes in HRV were noticed across 9 days of an international tournament (independently of being DT, DM, or EM). Regarding the training and match loads of the investigated soccer team, it appears that the s-RPE of training sessions and matches during the international tournament were not high enough to cause overnight changes in the cardiac autonomic system in this specific National team. Our study showed that, as a group, elite female soccer players are able to tolerate up to ≈3000 AU across 9 days or ≈600 AU per day of s-RPE during an international tournament, without presenting signs of severe nocturnal cardiac autonomic perturbation. Accordingly, Costa et al., 2018 [20] found in highly trained female soccer players during a 7-day period of the competitive season that late-night soccer training did not affect nocturnal HRV indices in comparison with rest days. The authors suggest that the late-night training loads, as measured by training impulse (range between: 77.5 [36.5] and 110.8 [31.6] AU) and s-RPE (range between: 281.8 [117.9] and 369.0 [111.7] AU), were not high enough to disturb the cardiac autonomic function during sleep hours. Although National team players might be more resilient to sustained high levels of training and match load without presenting signs of severe nocturnal cardiac autonomic perturbation, this needs to be further investigated.

### Paragraph 31

The lnRMSSD_CV_ has also been assessed in studies involving highly-trained athletes as a marker of weekly variation in daily assessed lnRMSSD [60]. In a recent study [61], the authors suggested that a high diurnal lnRMSSD_CV_ (subject 3 CV=12.8% and subject 8 CV=11.9%) was positively associated with perceived fatigue and negatively associated with the physical fitness of female soccer players. Moreover, another study found that diurnal lnRMSSD_CV_ measured in swimmers can increase to values>10% during overload periods [62]. In our study, as a group, lnRMSSD derived from the SWSE method displayed low average CV (6%). In fact, most of the lnRMSSD values across all days of the tournament were located within the SWC range (3%) [32]. However, individually, two players (players 6 and 7) appeared to present higher lnRMSSD_CV,_ which occurred simultaneously with a reduced average lnRMSSD during the tournament, contrasting with the remaining players of the team. Thus, overnight HRV during the international tournament was more sensitive to variation in these two players. Furthermore, it could be speculated that higher lnRMSSD_CV_ associated with reduced average lnRMSSD during training and matches may be interpreted as a sign of overload [61]. However, additional measures (e.g., well-being ratings) are needed to contextualize and better interpret changes in HRV [61, 63]. Unfortunately, due to time and technical constraints, well-being, fitness level, and other types of perceived stress were not determined, and we recognize this as an important limitation of the current study to better understand the HRV values of each player during the tournament.

### Paragraph 32

With the purpose of monitoring athletes, measurements performed during consecutive days throughout the week are very useful to interpret adaptations [64]. In our study, players seemed to display lower perturbation of nocturnal cardiac autonomic activity, as expressed by the lnRMSSD_CV_ [64]. Although the study comprised only 9 days of observation, it can be globally inferred that players were coping well with the training sessions and matches. In fact, a reduced lnRMSSD_CV_ without a clear decrease in the lnRMSSD, which occurred for most of these female soccer players, may reflect the possibility of a high level of readiness to perform [60] during the international tournament. This finding corroborates previous studies assessing HRV during the day, in athletes who were awake [26, 62]. Finally, coaches and technical staff should give great attention to player 6 who presented the highest CV for both TST and lnRMSSD (12.5% and 11.7%, respectively) during the 9 days of the international tournament.

### Paragraph 33. Limitations

Some limitations of the current study should be noted. Given that this study was conducted in a field setting (i.e., real-world scenario), there were many uncontrolled factors that may have affected the athletes’ sleep, other than match/training scheduling and load (e.g. use of caffeine; social media after lights out, electronic devices, level of light exposure during daytime, and perceived stress/well-being). In addition, caution should be applied when interpreting the CV values for sleep and HRV, since on some days there were missing data. Furthermore, this study was limited by the fact that a “real baseline” was not evaluated for possible sleep and nocturnal HRV comparisons across the 9 days of the tournament. Finally, while the session-RPE method may be simple, valid, and reliable, in this study we did not use other types of internal training and match load monitoring (e.g. HR monitors), which could provide objective information for the interpretation of the physiological responses of the players.

## Conclusions

### Paragraph 34

The present observational study is the first to systematically analyse consistent individual sleep and nocturnal HRV responses in elite female soccer players during a congested match schedule. Individually, some players presented less TST than recommended after some days of the tournament. However, the highest number of athletes sleeping less than 7 hours was found after EM compared with the remaining days of the tournament. Nevertheless, all players displayed good sleep quality for each day of the tournament. Additionally, most players seemed to present small fluctuations in nocturnal cardiac autonomic activity. Overall, elite female soccer players from a National team appeared to be highly resilient to training and match schedules and loads during an international tournament.

### Practical applications

#### Paragraph 35

Our observational analysis of sleep and nocturnal HRV responses of elite female soccer players could assist coaches and practitioners to identify sleep and HRV disturbances during official competitions, especially during periods of highly congested fixtures, as occurs during international tournaments. Moreover, this study highlights the substantial individual variability in sleep and HRV, suggesting the adoption of an individual approach to sleep (e.g. sleep hygiene), load monitoring, and recovery interventions in team sports.

## Acknowledgements

The authors would like to thank each of the athletes, and coaching and medical staff for their participation and cooperation during the study. FIFA Research Scholarship 2017 (International Centre for Sports Studies [CIES] and Fédération Internationale de Football Association [FIFA]) funding was provided for this study. The study was also supported by two individual doctoral grants from Fundação para a Ciência e a Tecnologia: Júlio Costa (SFRH/BD/128531/2017) and Vincenzo Rago (SFRH/BD/129324/2017).

